# A High-throughput Fluorescence Polarization Assay for Screening Sirtuin Inhibitors

**DOI:** 10.64898/2026.04.06.716694

**Authors:** Kewen Peng, Suryadeep Chakraborty, Yizhen Jin, Hening Lin

## Abstract

Sirtuins (SIRTs), which remove protein lysine acyl modifications, play crucial roles in diverse cellular processes, including metabolism, gene transcription, DNA damage repair, cell survival, and stress response. Several sirtuins are considered non-oncogene addiction of cancer cells and promising targets for anticancer drug development. High-throughput screening (HTS) methods for sirtuins are critical for the development of potent and isoform-selective sirtuin inhibitors, which are needed to validate the therapeutic potential. Herein, we designed and synthesized a fluorescent polarization (FP) tracer, KP-SC-1. Using this high-affinity tracer, we developed a robust, high-throughput FP competition assay for screening SIRT1-3 inhibitors. The assay was validated by testing known SIRT1-3 inhibitors. The assay can detect NAD^+^-independent SIRT1-3 inhibitors, as well as NAD^+^-dependent inhibitors, such as Ex-527 and TM. Finally, our assay showed satisfactory stability and outstanding performance in a pilot library screening. Compared to previous assays, the FP assay uses much less SIRT1-3 enzymes, a feature important for high-throughput library screening. We believe that the FP assay developed here will accelerate the discovery and development of SIRT1-3 inhibitors.

## INTRODUCTION

As class III histone deacetylases, sirtuins (SIRTs) remove acyl groups from protein lysine residues^1,2^. These enzymes use NAD^+^ as a co-substrate and transfer the acyl group from protein lysine residues to NAD^+^, producing O-acyl ADP-ribose, nicotinamide, and the deacylated protein. Humans have seven sirtuin proteins (SIRT1-SIRT7) that share a conserved core NAD^+^-binding catalytic domain, but they have distinct substrates, enzymatic activities, subcellular localizations, and functions^3,4^. Sirtuins have diverse biological roles, including metabolism, gene transcription, DNA damage repair, cell survival, and stress response^5–7^. Consequently, these enzymes have also been implicated in several human diseases, such as inflammation, neurodegenerative diseases, and cancers^8,9^. Among the seven human sirtuins, SIRT1, SIRT2, and SIRT3 belong to the same subfamily and have been studied extensively^5,7,10^.

Several sirtuins, including SIRT2, SIRT3, and SIRT5, are considered non-oncogene addiction of cancer cells. Thus, there have been sustained interest in developing sirtuin inhibitors^10–12^. Ex-527 is a SIRT1-selective inhibitor that reduced PANC-1 cell proliferation *in vitro*^13–15^. A mechanism-based SIRT2 inhibitor, TM, promotes the degradation of the oncoprotein c-Myc^16^. Subsequent structural modifications of TM led to other sirtuin inhibitors such as AF8^17^, YC8-02^18^ and SJ-106C,^19^ which exhibited distinct isoform selectivity and biological effects. Thieno[3,2-*d*]pyrimidine-6-carboxamide-based compound **11c** (Figure 1B) and several related compounds are pan-inhibitor of SIRT1-SIRT3 with nanomolar potency^20^.

**Figure 1.**
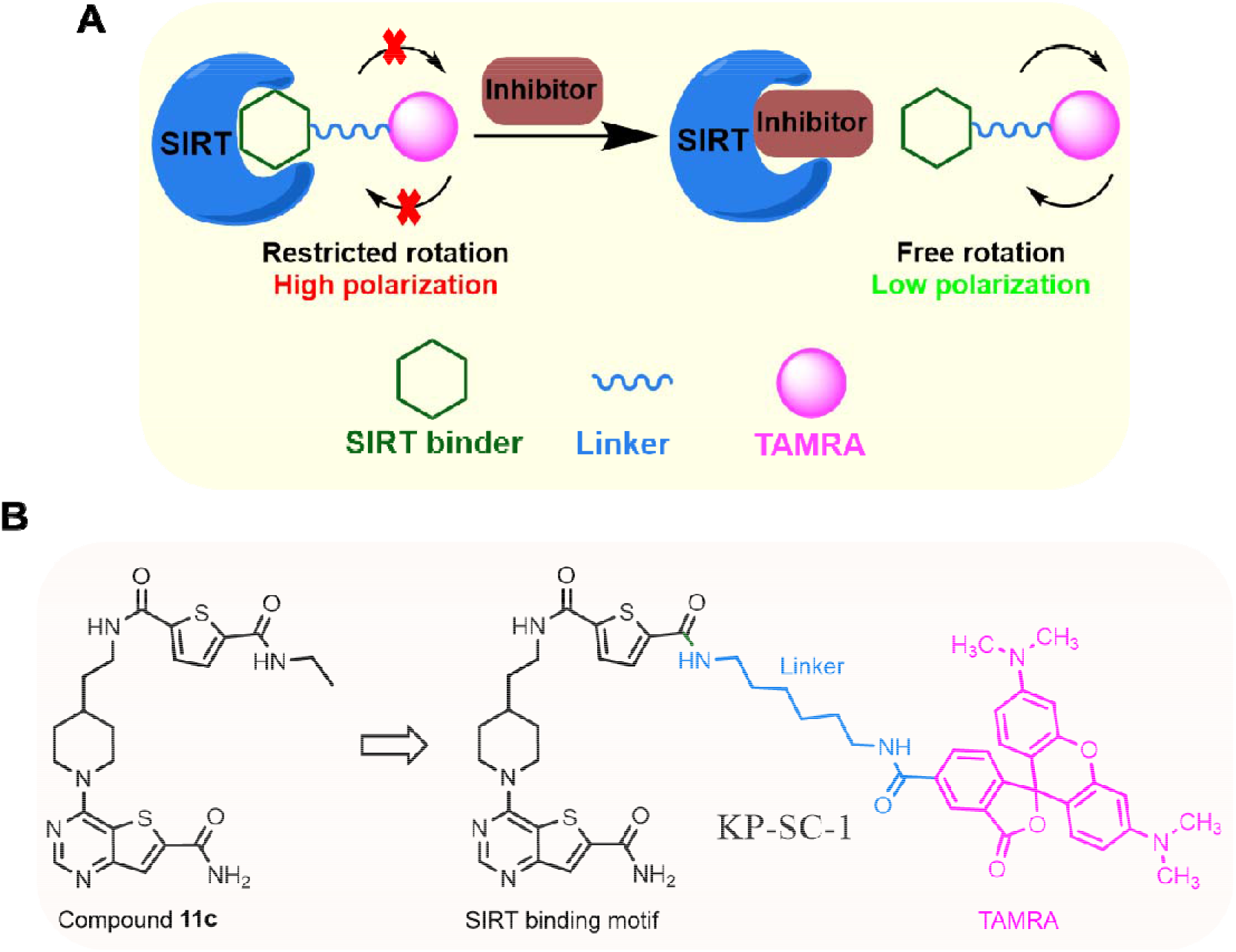
Mechanism and design of the fluorescence polarization (FP) assay for sirtuins. (A) When the FP tracer is bound to the sirtuin protein, its molecular rotation is restricted and thus generates high polarization. On the other hand, if the inhibitor replaces the tracer and thereby frees it from the protein, the tracer rotates rapidly, leading to low polarization. (B) Structure of FP tracer KP-SC-1. The fluorophore TAMRA (pink) is attached to the sirtuin-binding thieno[3,2-*d*]pyrimidine-6-carboxamide scaffold (black) via a diamide linker (blue).

Despite these efforts in sirtuin inhibitor development, current sirtuin inhibitors still need improvement^21,22^ due to various limitations. For instance, in contrast to its *in vitro* anticancer effect, Ex-527 promotes PANC-1 xenograft tumor growth *in vivo* in SCID mice^15^. The SIRT2-selective inhibitor TM has poor drug-like properties,such as low solubility^23^. On the other hand, the SIRT3 inhibitor SJ-106C, albeit being more soluble, does not have satisfactory SIRT3 selectivity^19^. Therefore, there is still a strong need to develop better sirtuin inhibitors.

The need to develop better sirtuin inhibitors underscores the importance of high-throughput screening (HTS) assays so that novel chemical entities targeting sirtuins can be quickly identified and subsequently optimized. Over the past few decades, several assays for sirtuins have been developed, including high-performance liquid chromatography (HPLC)-based assays, radioisotope-labelled ^32^P-NAD^+^ assays, mass spectrometry (MS)-based assays, and fluorogenic assays^24–26^. Among these, the fluorogenic assays are best suited for HTS. Fluorogenic assays can be further divided into direct fluorescence assays and fluorescence resonance energy transfer (FRET)-based assays^24,27,28^. In a direct fluorescence assay, 7-amino-4-methylcoumarine (AMC) is attached to the C-terminus of the substrate peptide comprising *ε*-acylated lysine residue, while in a FRET-based assay, the FRET pairs DABCYL-EDANS are coupled to the N-terminus and C-terminus of the acyl lysine substrate peptide, respectively. In both cases, the SIRT enzyme first removes the acyl group from the lysine residue. Subsequent trypsin-catalyzed digestion of the deacylated peptide either releases the AMC fluorophore in a direct fluorescence assay or disrupts the FRET effects in a FRET-based assay. These fluorogenic assays have several limitations. First, since trypsin digestion is indispensable, compounds that inhibit trypsin activity will show up as false positives. Furthermore, the two-step procedures of these assays render them more time-consuming and labor-intensive for large-scale screening applications. Finally, nearly all these reported assays, whether fluorogenic or not, require protein concentrations in the micromolar range. This high protein concentration requirement adds to the cost of screening campaigns.

The fluorescence polarization (FP) assay is a popular binding assay with a simple mix-and-read procedure, and has been widely used in HTS. The FP assay relies on the fact that when a fluorescent tracer (i.e., a fluorophore-conjugated ligand) binds to its partner protein, its freedom of molecular rotation is restricted, resulting in high polarization (Figure 1A)^29,30^. Therefore, the FP assay can be very cheap as only a tracer and the protein of interest are required.

Herein, we designed and synthesized a novel FP tracer KP-SC-1 with nanomolar binding affinity for sirtuins. With this tracer in hand, we established a stable FP assay with a wide assay window and successfully applied it to SIRT1, SIRT2, and SIRT3. Using our assay, we validated the binding affinities of inhibitors reported in the literature. In addition to detecting direct sirtuin binders, our assay also works for NAD^+^-dependent sirtuin inhibitors such as Ex-527 and TM. Importantly, by conducting the assay in the presence or absence of NAD^+^, we can differentiate NAD^+^-dependent and independent inhibitors. Finally, we employed the assay to screen a fragment library and demonstrated its outstanding performance with satisfactory %CV and Z^′^ factor.

## RESULTS and DISCUSSION

**Figure Sc1.**
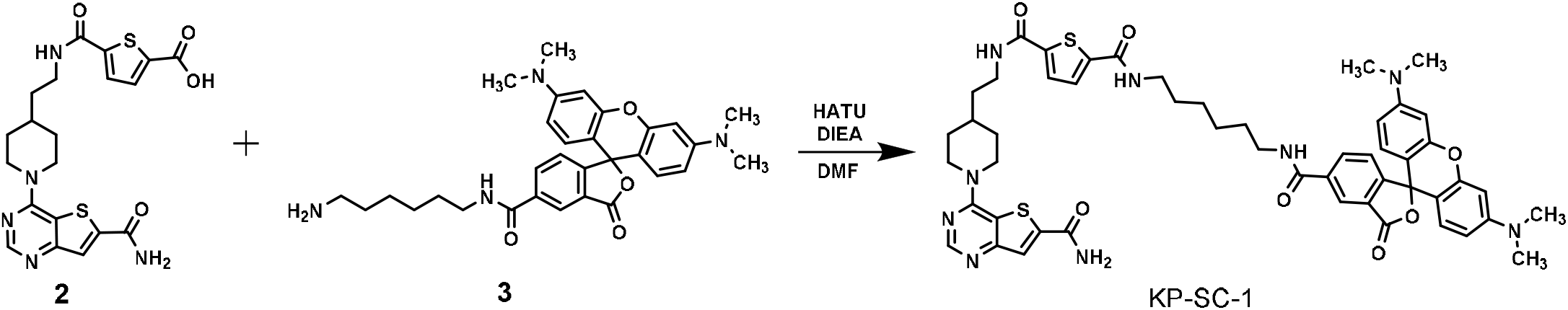
Synthetic route for KP-SC-1

Compound **11c**, a thieno[3,2-*d*]pyrimidine-6-carboxamide derivative, has been reported to inhibit SIRT1, SIRT2, and SIRT3 with nanomolar affinities^20^, and thus represent a promising starting point for a SIRT1-3 tracer design (Figure 1B). Since the terminal N-ethyl amide position of compound **11c** can tolerate substitutions as bulky as DNA tags, we envisioned that this site should be suitable for the installation of a fluorescent tag. We replaced the N-ethyl amide group with a six-carbon diamide-based linker that connects to a bright purple fluorophore, TAMRA, and designed compound KP-SC-1 (Figure 1B). We synthesized KP-SC-1 following the synthetic route depicted in Scheme 1.

With tracer KP-SC-1 in hand, we next examined its binding affinity for sirtuins through FP titration. In this titration assay, a fixed concentration of tracer (2 nM) was titrated with increasing concentrations of sirtuins (SIRT1-3). The mP shift (ΔmP) was calculated as the difference between the mP values of the protein-tracer sample and the tracer-only sample. The binding curves (Figure 2A) indicated a strong affinity of KP-SC-1 for all three sirtuin proteins. The *K*_d_ values of KP-SC-1 were calculated to be 32 nM, 6 nM, and 10 nM for SIRT1, SIRT2, and SIRT3, respectively. This is consistent with the reported activity data of compound **11c**^20^. We also tested other sirtuins like SIRT5-7 in the FP assay. However, the tracer did not bind to these sirtuins (Figure S1).

**Figure 2.**
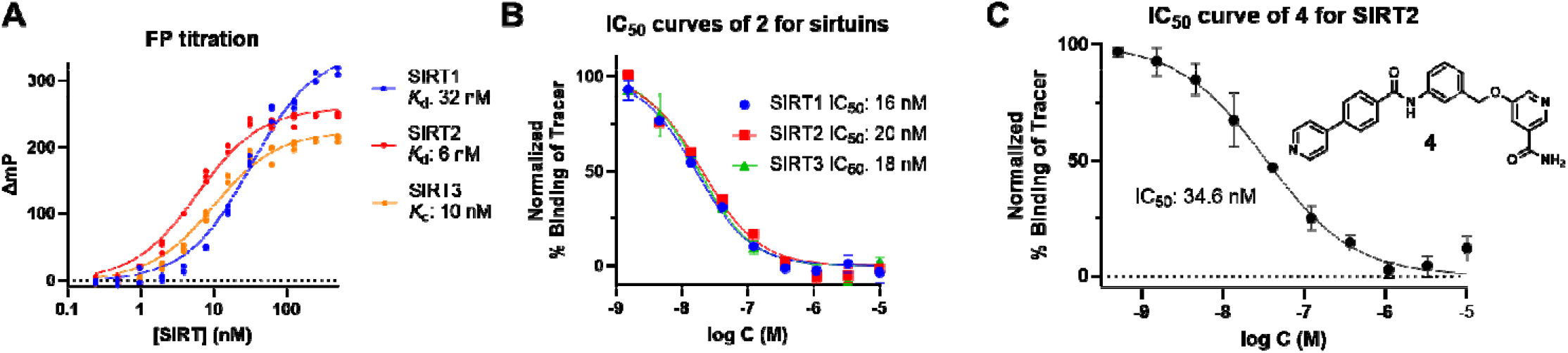
FP titration and assay validation. (A) FP titration of 2 nM of tracer KP-SC-**1** against SIRT1, SIRT2, and SIRT3 (n=3). (B) Using sirtuin inhibitor **2** to validate the FP assay with the following condition: 30 nM SIRT1, 10 nM SIRT2, or 15 nM SIRT3 with 2 nM tracer in 100 μL final volume in 96-well plates (n=2). (C) Using SIRT2 inhibitor **4** to validate the FP assay with the following condition: 25 nM SIRT2 and 5 nM tracer in 40 μL final volume in a 384-well plate (n=4). The structure of **4** is also shown.

The optimal concentrations of tracer and protein were selected to ensure sufficient fluorescence signal and an mP window of 150. In general, the protein was used at around 1- to 3-fold *K*_d_, and the tracer was used at 2-5 nM in 96-well plates or 5-10 nM in 384-well plates. To validate the assay, we measured the binding affinity of **2** for SIRT1-3. Different concentrations of **2** were incubated with a fixed concentration of sirtuin for 30 min at room temperature, and the parallel and perpendicular fluorescence intensities were then measured with a plate reader. Subsequently, the inhibitor concentrations in log unit were plotted against the normalized percentage of binding of tracer to generate IC_50_ curves (Figure 2B). The IC_50_ values obtained for **2** were 16 nM, 20 nM, and 18 nM for SIRT1, SIRT2, and SIRT3, respectively, which are in line with reported values of 14 nM, 4.4 nM, and 13 nM for this compound^20^. Compound **4**,^31^ a potent and selective SIRT2 inhibitor, was chosen as another test case for assay validation. The IC_50_ of **4** in our assay was calculated to be 34.6 nM (Figure 2C), which is consistent with the reported IC_50_ of 22.9 nM measured in an AMC-based fluorogenic assay.

High assay stability is desirable for library screening purposes, since it takes considerably longer time to read the plates in large-scale library screening. Most plate readers can only read one well at a time, and particularly in FP assays, the plate must be read twice for both parallel and perpendicular fluorescence values.Therefore, data homogeneity is important during the time frame when the plates are being read, whether that being minutes or hours. To interrogate our assay stability over time for SIRT1-3, we incubated the assay plate for 6 hours at room temperature and monitored the mP values at different time points. Ideally, mP should remain unchanged when the protein and/or tracer are not degraded, and their binding interaction is maintained. Indeed, the mP values for SIRT2 and SIRT3 remained unchanged over 6 hours (Figure 3B and 3C). For SIRT1, the mP decreased gradually over time (Figure 3A). We found that under the conditions we used, SIRT1 formed higher molecular weight oligomer (Figure S3). The oligomer formation was prevented by the addition of DTT, suggesting that the oligomer formation is likely mediated by disulfide bonds. Accordingly, the presence of DTT in SIRT1 FP prevented the mP drop (Figure S4). Therefore, we recommend adding DTT when conducting SIRT1 FP assay.

**Figure 3.**
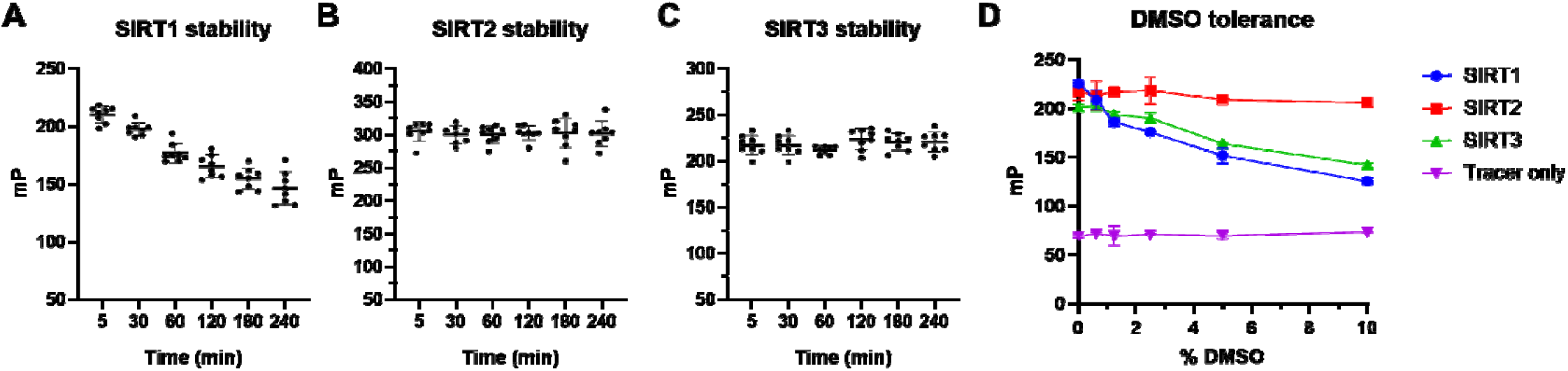
FP assay stability over time for (A) SIRT1, (B) SIRT2, and (C) SIRT3 with tracer KP-SC-1. SIRT1, 2, and 3 were used at 60 nM, 30 nM, and 30 nM, respectively. Tracer KP-SC-1 concentration was 5 nM, and the assay volume was 40 μL in a 384-well plate (n=8). (D) Effect of different percentages of DMSO on mP values of tracer KP-SC-1 with or without SIRT1/2/3 (n=2).

Since DMSO can affect protein structure and affect protein-tracer binding, DMSO tolerance of the assay is another important factor to consider, as in many cases high DMSO may be needed for testing weaker hits.Therefore, we examined the DMSO tolerance of our assay by conducting the assay in buffers with different DMSO contents. As shown in Figure 3D, SIRT1 was very sensitive to DMSO, and its mP values decreased as DMSO content was increased. However, SIRT3 could tolerate up to 2.5% DMSO with minimal effects on the mP value. Strikingly, even 10% DMSO did not affect the stability of SIRT2. Our data suggest SIRTs have distinct DMSO stability profiles, and DMSO contents should be used at < 1% for SIRT1.

Some sirtuin inhibitors, such as Ex-527, TM, and SJ-106C, require NAD^+^ for their inhibitory effect. The binding of Ex-527 to SIRT1 requires NAD^+14^. Thioamide-containing inhibitors, such as TM and SJ-106C, form a stalled covalent intermediate with NAD^+^ to inhibit sirtuins^16,19^. As our developed FP tracer directly binds NAD^+^ binding pockets, our assay in its original form might not identify these NAD^+^-dependent inhibitors.Indeed, when we tested Ex-527 and TM against SIRT1 and SIRT2, respectively, no inhibitory effect was observed for either compound in the absence of NAD^+^ (Figure S2A and S2B).

To extend our FP assay’s applicability to NAD^+^-dependent inhibitors, we sought to include NAD^+^ in the assay. One major concern of adding NAD^+^ is that NAD^+^ itself may compete with our tracer and thus significantly decrease the assay window. Gratifyingly, we did not observe any decrease in mP values for all SIRTs when NAD^+^ was used at 1 mM (Figure 4A), a concentration commonly used in SIRT enzyme assays. This observation, while initially surprising, actually fits the known sirtuin enzymology literature. All human sirtuins, except SIRT6, bind NAD^+^ only in the presence of acyl peptide substrate^32–35^. In our FP assay, there is no acyl peptide present and thus, NAD^+^ has very weak binding affinities towards SIRT1, SIRT2, and SIRT3,and does not interfere with the binding of the FP probe.

**Figure 4.**
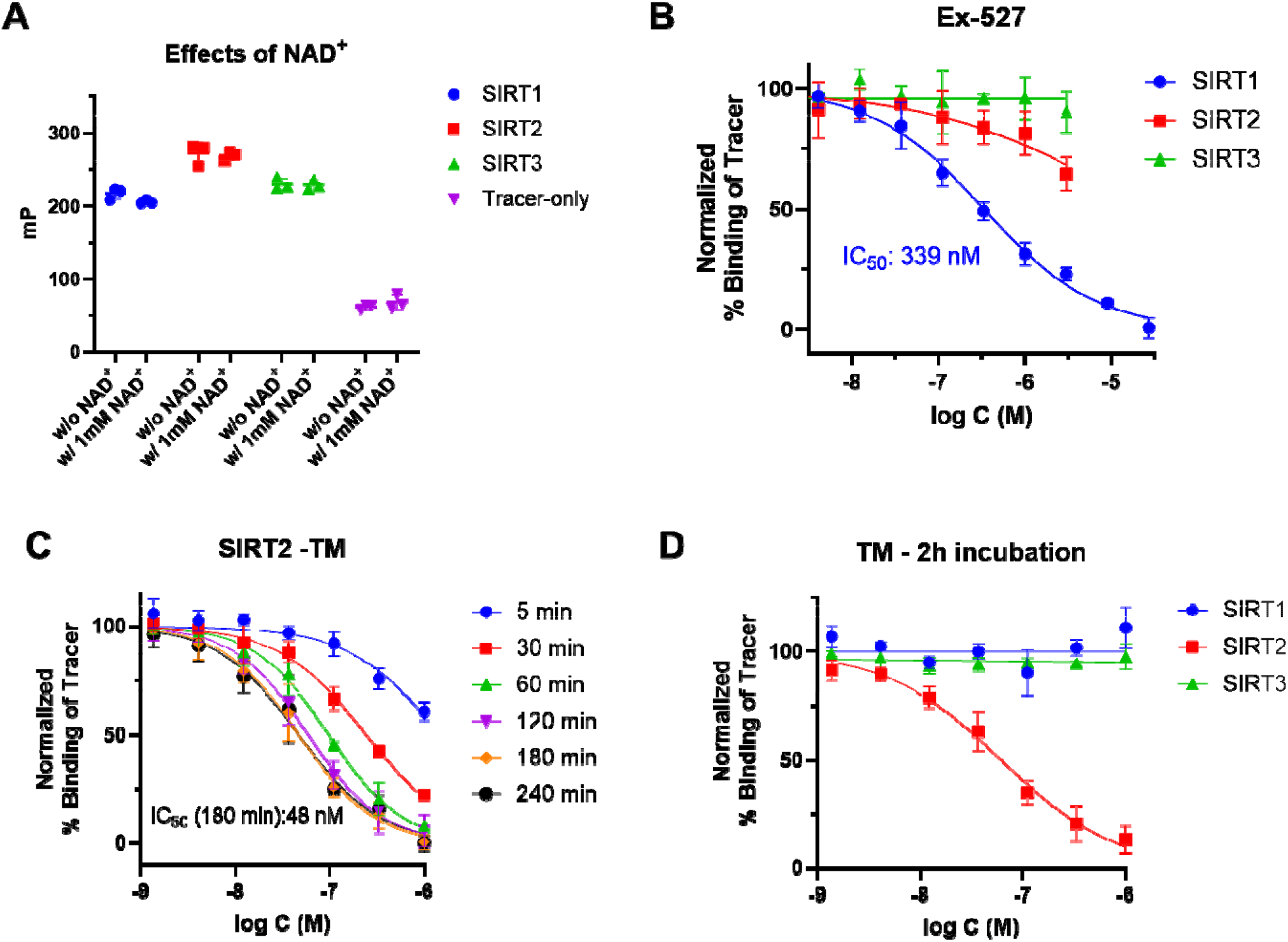
The FP assay is applicable to NAD^+^-dependent inhibitors. (A) NAD^+^ at 1 mM has no significant effects on the mP values (n=3); (B) IC_50_ curves of the SIRT1-selective inhibitor Ex-527 (n=4); (C) IC_50_ curve of SIRT2 inhibitor TM with different incubation time. IC_50_ value of TM with 180 min incubation time is shown (n=4); (D) IC_50_ curves of TM with SIRT1, SIRT2, and SIRT3. TM shows selective SIRT2 inhibition (n=4).

Subsequently, we examined the inhibitory effects of the SIRT1-selective Ex-527 and the SIRT2-selective TM in our FP assay in the presence of 1 mM NAD^+^. Ex-527 exhibited an IC_50_ of 339 nM for SIRT1 (Figure 4B), which is close to the reported IC_50_ of ∼98 nM obtained from enzyme assays^13^, and did not inhibit SIRT2 or SIRT3. Similarly, TM inhibited SIRT2 but not SIRT1 or SIRT3 as expected (Figure 4D). Interestingly, the inhibitory effect of TM increased with the incubation time (Figure 4C), which suggests TM requires sufficient time to form the active adduct with NAD^+^. At 3 h, TM showed an IC_50_ of 48 nM for SIRT2, which is consistent with the reported value of 28 nM^16^. Therefore, another advantage of our FP assay is that NAD^+^ can be optionally decoupled from the assay to interrogate compound’s inhibition mechanism, which is imposible to achieve in traditional enzyme assays where NAD^+^ is indispensable.

To test the performance of our FP assay in a library screening setting, we performed a pilot screen of the Enamine Essential Fragment Library comprising 320 fragments, using SIRT3 as the test protein (Figure 5A).Six fragments with intrinsic fluorescence were excluded. The library was screened at 100 µM, with 10 nM tracer and 30 nM SIRT3 in each well in a 384-well plate. The tracer and SIRT3 concentrations used were much lower than the enzyme and substrate concentrations used in previous fluorogenic assays, which is a very desirable feature for HTS. Appropriate controls, including buffer control, negative control (protein + tracer), and positive control (only tracer), were used. From the controls, we calculated the assay window, ΔmP, which was more than 100 (Figure 5B), higher than the commonly accepted threshold of 70^36^. The screening window coefficient (Z^′^ factor), which evaluates the robustness and quality of a HTS assay, was calculated to be 0.79 (Figure 5C), falling into the desirable range between 0.5 and 1^37^. Additionally, the percentage coefficient of variation (% CV) is low for both negative (3.6 %) and positive controls (2.7%), indicating good reproducibility and precision of our assay. Although our pilot library screening did not yield any hit compounds, it demonstrated the assay’s superior performance in HTS, setting up the stage for future screening of more sophisticated compound libraries.

**Figure 5.**
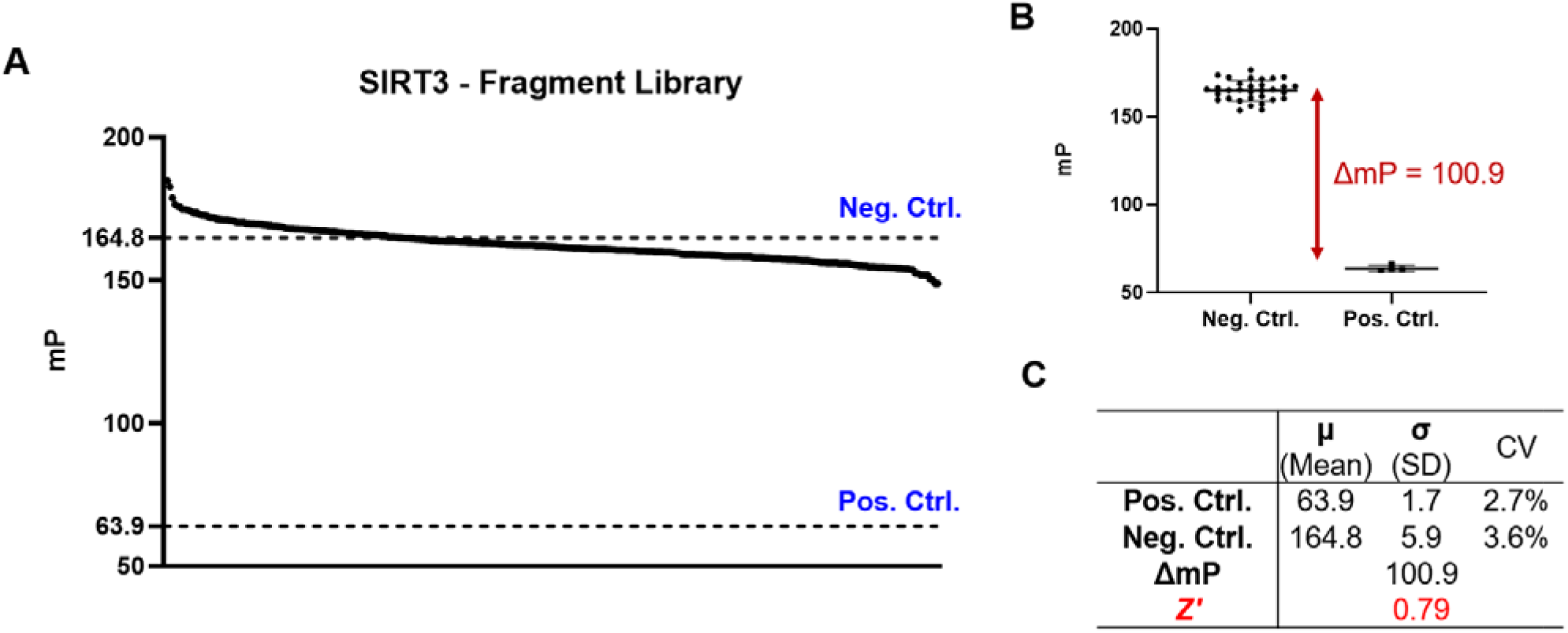
Pilot screening of the Enamine Essential Fragment library using the FP assay with SIRT3. SIRT3 and tracer concentration were 30 nM and 10 nM, respectively. The screening was performed in a 384-well plate at a volume of 40 µL for each well (n=1). (A) Data from the fragment screening. The negative (SIRT3 and tracer only) and positive (tracer only) controls are shown with the horizontal dotted lines. (B) Negative and positive control data. The assay window () was calculated as the difference between the means of the negative and positive controls. (C) Summary of key screening parameters, %CV,, and Z′.

## CONCLUSION

We developed a fluorescent tracer, KP-SC-1, and developed an FP-based competition assay for screening SIRT1-3 inhibitors. We demonstrated that the tracer has nanomolar binding affinity for SIRT1-3. The tracer does not work for other sirtuin isoforms like SIRT5-7, probably due to significant differences in their active sites. The assay was validated using known sirtuin inhibitors. The assay shows high temporal and DMSO stability for SIRT2 and SIRT3, whereas for SIRT1, its stability is less desirable but still acceptable for general screens. We further included NAD^+^ in the assay and showed that it could also detect NAD^+^-dependent inhibitors. Finally, we showed that the FP assay performed well in high-throughput screening. This assay will be helpful in developing selective and potent sirtuin inhibitors (both NAD^+^-dependent and NAD^+^-independent) in a high-throughput manner.

## MATERIALS and METHODS

### Reagents

NAD^+^ (SKU: N30110-1.0) was obtained from RPI (RESEARCH PRODUCTS International). Ex-527 (Cat No. S1541) was purchased from Selleck Chemicals LLC. The Enamine Fragment Library (Catalog Number: ESS-320-100-X-100) was purchased from Enamine. Recombinant human N-terminal His-tagged SIRT6 (Item No. 10315) and recombinant human C-terminal FLAG-tagged SIRT7 (Catalog Number: 50018) were purchased from CAYMAN CHEMICAL and BPS Bioscience, respectively. Unless otherwise noted, all other biological reagents and consumables were obtained from commercial vendors.

### Chemical Synthesis

The details of the chemical synthesis can be found in the supporting information.

### Cloning, Expression, and Purification of Human Sirtuins

Human SIRT1, SIRT2, SIRT3, and SIRT5 were expressed and purified as previously described^16^.

### FP Titrations of sirtuins

Using the assay buffer (25 mM Tris, pH 8.0, 150 mM NaCl, 0.01% Tween-20), the purified SIRT1-SIRT3 proteins were serially diluted 2-fold from 1000 nM to ∼ 0.5 nM. The assay was performed in a 96-well flat-bottom black microplate (Corning Product Number: 3915). The titration assay was performed for all three SIRT1, SIRT2, and SIRT3 following the same protocol. For the titration assay, 50 µL of each protein concentration (2X) was transferred to separate wells of the microplate. Subsequently, 50 µL of FP tracer KP-SC-1 (4 nM, 2X) was added to each well, bringing the total volume to 100 µL per well. Mixing equal volumes of protein and tracer in each well resulted in a final concentration of 1X. The plate was incubated at room temperature for 30 min and then scanned on a Cytation 5 equipped with a filter cube (Agilent, part number: 8040562; Ex: 530/25, Em: 590/35, Cut-off: 570). Fluorescence intensities parallel and perpendicular to the plane of the linearly polarized excitation light were measured. Then the blank-subtracted fluorescence intensities were used to calculate the mP values using the following equation^30^

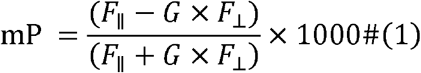

where *F*_∥_ and *F*_⊥_ are parallel and perpendicular fluorescence intensities, respectively, and G is the instrument’s grating factor. Subsequently, the ΔmP was calculated for each well by subtracting the mP value of the tracer-only control from the mP of the corresponding sample. The resulting ΔmP values were fitted to the one-site specific binding model implemented in GraphPad Prism 10.6.1 (GraphPad Software, Inc.), and *K*_d_ values were calculated using the following equation

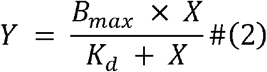

Where *X* and *Y* represent the protein concentration and binding response (mP shift), respectively. *B*_*max*_ is the maximum binding response.

### FP-based Binding Assay for sirtuins with the Inhibitors

The stock solution of sirtuins and the tracer KP-SC-1 were diluted with the assay buffer (25 mM Tris, pH 8.0, 150 mM NaCl, 0.01% Tween-20) to a 4X concentration. Tracer KP-SC-1 (4X concentration) was added to an equal volume of protein solution to make a protein-tracer solution of 2X concentration for each component. Using the assay buffer, sirtuin inhibitors were serially diluted. These serially diluted inhibitor solutions (2X concentrations) were transferred to a 96-well microplate at 50 µL per well. Subsequently, 50 µL of the protein-tracer mixture (2X concentration) was added to the wells containing the inhibitors. The final concentrations of each component (protein, tracer, and inhibitors) become 1X in the assay. To test the inhibitory effect of Ex-527 or TM in the presence of 1 mM NAD^+^, all dilutions were performed with a 1 mM NAD^+^ stock solution prepared in the assay buffer. In the cases where the binding assay was performed in a 384-well plate, the total assay volume was kept at 40 µL, and the other procedures were followed in the same manner as mentioned above. Controls include the negative control (the inhibitor was replaced with buffer), the tracer control (2X concentration of tracer was mixed with buffer instead of protein), and the buffer control (no protein or tracer). The percentage binding of tracer relative to the controls was calculated using the following equation

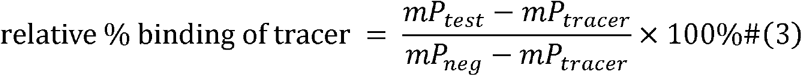

Where *mP*_*test*_, *mP*_*tracer*_ and *mP*_*neg*_ represent the mP values of the test wells, the tracer-only control, and the negative control, respectively. The data were plotted in GraphPad Prism 10.6.1 (GraphPad Software, Inc.), with the X-axis as log C (M) and the Y-axis as the normalized % binding of tracer. The data were fitted to an IC_50_ curve using the sigmoidal four-parameter logistic model, using the following equation:

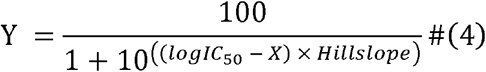

where X and Y are the inhibitor concentration and the relative % of binding of tracer, respectively. Statistical values including mean, standard deviation (SD) and coefficient of variation (CV) were obtained with Excel functions.

### Stability and DMSO Tolerance of the Assay

Stock solutions of sirtuins and tracer KP-SC-1 were separately diluted in assay buffer (25 mM Tris, pH 8.0, 150 mM NaCl, 0.01% Tween-20) to 2X concentration. Equal volumes of each were mixed on a 384-well black plate (Corning Product Number: 3575) to make a final concentration of 1X. The final volume was maintained at 40 µL (20 µL protein + 20 µL tracer). The final concentrations of tracer, SIRT1, SIRT2, and SIRT3 were 5 nM, 60 nM, 30 nM, and 30 nM, respectively. The fluorescence intensities of the sample were measured at 5 min, 30 min, 1 h, 2 h, 3 h, and 4 h. Subsequently, the fluorescence data were converted to mP data and plotted in GraphPad Prism 10.6.1 (GraphPad Software, Inc.) over different time courses in the X-axis.

For the DMSO tolerance assay, two solutions of tracer were prepared: 10 nM tracer KP-SC-1 in 20 % (volume-wise) DMSO, and 10 nM tracer KP-SC-1 in assay buffer. Serial dilution of DMSO was made by mixing an equal volume of tracer KP-SC-1 in DMSO and 10 nM tracer KP-SC-1 in assay buffer. Thus, this equal-volume mixing will keep the tracer concentration at 10 nM while diluting the DMSO by 2-fold. Following this procedure, 10 nM tracer was prepared in different DMSO volume percentages, namely 20 %, 10 %, 5 %, 2.5 %, and 1.25 %. Subsequently, the tracer solutions with different DMSO concentrations were transferred into a 384-well black plate (Corning Product Number: 3575), with 20 µL in each well. Subsequently, 20 µL of protein stock (2X concentration) was added to the wells with tracer. The final concentrations of tracer, SIRT1, SIRT2, and SIRT3 were 5 nM, 60 nM, 30 nM, and 40 nM, respectively. The final DMSO concentrations were 10%, 5%, 2.5%, 1.25%, 0.625%, and 0%. Only tracer (i.e., without any protein) was also incubated with the same DMSO concentration as a control.

### Fragment Library Screening

39.6 µL of the assay solution (30 nM SIRT3 and 10 nM tracer KP-SC-1) or control solution (tracer only, buffer only) was transferred from a 96-well mother plate to a 384-well black plate (Corning Product Number: 3575). Subsequently, 0.4 µL of 10 mM fragment in DMSO was added to each well except the control wells, where 0.4 µL of DMSO was added. The plate was incubated for 30 min before being scanned as mentioned above. For each fragment, positive and negative control, the relative percentage of binding of tracer was calculated as described above. Z^′^ was determined using the following equation

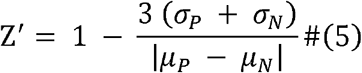

where *µ*_*P*_ and *µ*_*N*_ are the mean ΔmP values of the positive and negative control, respectively. *σ*_*P*_ and,*σ*_*N*_ represent the standard deviation of ΔmP values of the positive and negative control, respectively.

## Supporting information

Supplementary Figures and synthesis procedure

## Associated Content

### Supporting Information

Chemical synthesis and analytical data for the probe; additional experimental data (Figure S1-S4).

## Acknowledgement

The work is supported in part by NIH/NCI grant R01CA270243. The NMR spectra was obtained with assistance from the University of Chicago NMR Facility. HRMS data was obtained in the University of Chicago Mass Spectrometry Facility, which is supported by the NSF instrumentation grant CHE-1048528.

## Author Contributions

K.P. designed and synthesized the probe. K.P., S.C., and Y.J. performed the biochemical experiments. K.P. and S.C. analyzed the data. S.C. wrote the manuscript with revisions made by all authors. H.L. conceptualized and supervised this project.

## Conflict of Interest

The authors declare no conflict of interest.

---

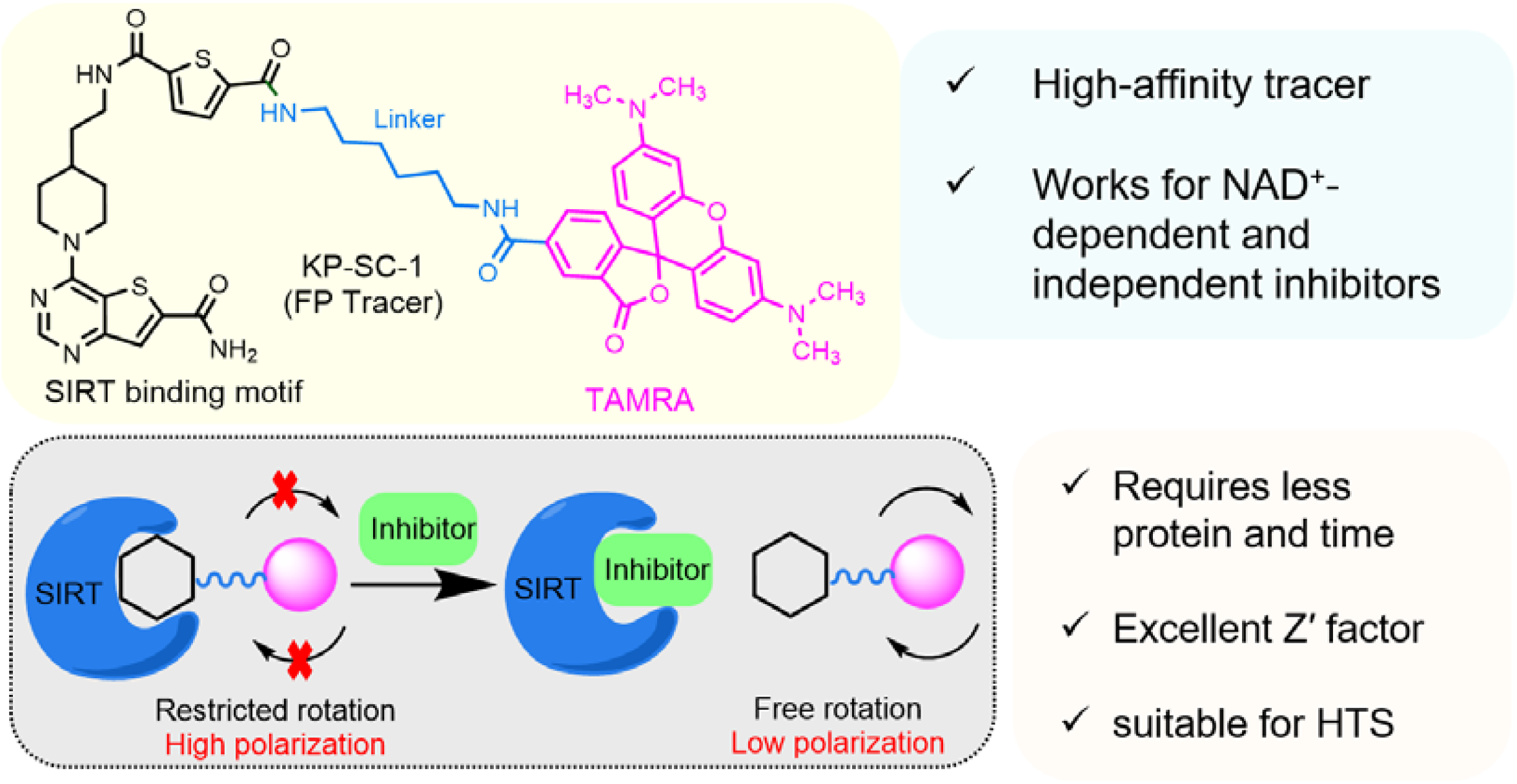

## Notes

### Competing Interest Statement

The authors have declared no competing interest.

### Summary of Updates

SIRT1 assay stability (Figure S3 and S4); discussion about why NAD+ does not affect the probe binding.

## References

(1) Wang, M.; Lin, H. Understanding the Function of Mammalian Sirtuins and Protein Lysine Acylation. Annu Rev Biochem 2021, 90, 245–285. 10.1146/annurev-biochem-082520-125411.

(2) Sauve, A. A.; Wolberger, C.; Schramm, V. L.; Boeke, J. D. The Biochemistry of Sirtuins. Annu. Rev. Biochem. 2006, 75 (1), 435–465. 10.1146/annurev.biochem.74.082803.133500.

(3) Houtkooper, R. H.; Pirinen, E.; Auwerx, J. Sirtuins as Regulators of Metabolism and Healthspan. Nat Rev Mol Cell Biol 2012, 13 (4), 225–238. 10.1038/nrm3293.

(4) Bheda, P.; Jing, H.; Wolberger, C.; Lin, H. The Substrate Specificity of Sirtuins. Annu. Rev. Biochem. 2016, 85 (1), 405–429. 10.1146/annurev-biochem-060815-014537.

(5) Jing, H.; Lin, H. Sirtuins in Epigenetic Regulation. Chem Rev 2015, 115 (6), 2350–2375. 10.1021/cr500457h.

(6) Yu, L.; Li, Y.; Song, S.; Zhang, Y.; Wang, Y.; Wang, H.; Yang, Z.; Wang, Y. The Dual Role of Sirtuins in Cancer: Biological Functions and Implications. Front. Oncol. 2024, 14, 1384928. 10.3389/fonc.2024.1384928.

(7) Kosciuk, T.; Wang, M.; Hong, J. Y.; Lin, H. Updates on the Epigenetic Roles of Sirtuins. Current Opinion in Chemical Biology 2019, 51, 18–29. 10.1016/j.cbpa.2019.01.023.

(8) Wu, Q.-J.; Zhang, T.-N.; Chen, H.-H.; Yu, X.-F.; Lv, J.-L.; Liu, Y.-Y.; Liu, Y.-S.; Zheng, G.; Zhao, J.-Q.; Wei, Y.-F.; Guo, J.-Y.; Liu, F.-H.; Chang, Q.; Zhang, Y.-X.; Liu, C.-G.; Zhao, Y.-H. The Sirtuin Family in Health and Disease. Signal Transduct Target Ther 2022, 7 (1), 402. 10.1038/s41392-022-01257-8.

(9) Herskovits, A. Z.; Guarente, L. Sirtuin Deacetylases in Neurodegenerative Diseases of Aging. Cell Res 2013, 23 (6), 746–758. 10.1038/cr.2013.70.

(10) Hu, J.; Jing, H.; Lin, H. Sirtuin Inhibitors as Anticancer Agents. Future Med. Chem. 2014, 6 (8), 945–966. 10.4155/fmc.14.44.

(11) Chalkiadaki, A.; Guarente, L. The Multifaceted Functions of Sirtuins in Cancer. Nat Rev Cancer 2015, 15 (10), 608–624. 10.1038/nrc3985.

(12) Jeong, S. M.; Haigis, M. C. Sirtuins in Cancer: A Balancing Act between Genome Stability and Metabolism. Molecules and Cells 2015, 38 (9), 750–758. 10.14348/molcells.2015.0167.

(13) Napper, A. D.; Hixon, J.; McDonagh, T.; Keavey, K.; Pons, J.-F.; Barker, J.; Yau, W. T.; Amouzegh, P.; Flegg, A.; Hamelin, E.; Thomas, R. J.; Kates, M.; Jones, S.; Navia, M. A.; Saunders, J. O.; DiStefano, P. S.; Curtis, R. Discovery of Indoles as Potent and Selective Inhibitors of the Deacetylase SIRT1. J Med Chem 2005, 48 (25), 8045–8054. 10.1021/jm050522v.

(14) Gertz, M.; Fischer, F.; Nguyen, G. T. T.; Lakshminarasimhan, M.; Schutkowski, M.; Weyand, M.; Steegborn, C. Ex-527 Inhibits Sirtuins by Exploiting Their Unique NAD^+^ -Dependent Deacetylation Mechanism. Proc. Natl. Acad. Sci. U.S.A. 2013, 110 (30). 10.1073/pnas.1303628110.

(15) Oon, C. E.; Strell, C.; Yeong, K. Y.; Östman, A.; Prakash, J. SIRT1 Inhibition in Pancreatic Cancer Models: Contrasting Effects in Vitro and in Vivo. Eur J Pharmacol 2015, 757, 59–67. 10.1016/j.ejphar.2015.03.064.

(16) Jing, H.; Hu, J.; He, B.; Negrón Abril, Y. L.; Stupinski, J.; Weiser, K.; Carbonaro, M.; Chiang, Y.-L.; Southard, T.; Giannakakou, P.; Weiss, R. S.; Lin, H. A SIRT2-Selective Inhibitor Promotes c-Myc Oncoprotein Degradation and Exhibits Broad Anticancer Activity. Cancer Cell 2016, 29 (3), 297–310. 10.1016/j.ccell.2016.02.007.

(17) Farooqi, A. S.; Hong, J. Y.; Cao, J.; Lu, X.; Price, I. R.; Zhao, Q.; Kosciuk, T.; Yang, M.; Bai, J. J.; Lin, H. Novel Lysine-Based Thioureas as Mechanism-Based Inhibitors of Sirtuin 2 (SIRT2) with Anticancer Activity in a Colorectal Cancer Murine Model. J. Med. Chem. 2019, 62 (8), 4131–4141. 10.1021/acs.jmedchem.9b00191.

(18) Li, M.; Chiang, Y.-L.; Lyssiotis, C. A.; Teater, M. R.; Hong, J. Y.; Shen, H.; Wang, L.; Hu, J.; Jing, H.; Chen, Z.; Jain, N.; Duy, C.; Mistry, S. J.; Cerchietti, L.; Cross, J. R.; Cantley, L. C.; Green, M. R.; Lin, H.; Melnick, A. M. Non-Oncogene Addiction to SIRT3 Plays a Critical Role in Lymphomagenesis. Cancer Cell 2019, 35 (6), 916-931.e9. 10.1016/j.ccell.2019.05.002.

(19) Jana, S.; Shang, J.; Hong, J. Y.; Fenwick, M. K.; Puri, R.; Lu, X.; Melnick, A. M.; Li, M.; Lin, H. A Mitochondria-Targeting SIRT3 Inhibitor with Activity against Diffuse Large B Cell Lymphoma. J Med Chem 2024, 67 (17), 15428–15437. 10.1021/acs.jmedchem.4c01053.

(20) Disch, J. S.; Evindar, G.; Chiu, C. H.; Blum, C. A.; Dai, H.; Jin, L.; Schuman, E.; Lind, K. E.; Belyanskaya, S. L.; Deng, J.; Coppo, F.; Aquilani, L.; Graybill, T. L.; Cuozzo, J. W.; Lavu, S.; Mao, C.; Vlasuk, G. P.; Perni, R. B. Discovery of Thieno[3,2-d]Pyrimidine-6-Carboxamides as Potent Inhibitors of SIRT1, SIRT2, and SIRT3. J Med Chem 2013, 56 (9), 3666–3679. 10.1021/jm400204k.

(21) Dai, H.; Sinclair, D. A.; Ellis, J. L.; Steegborn, C. Sirtuin Activators and Inhibitors: Promises, Achievements, and Challenges. Pharmacology & Therapeutics 2018, 188, 140–154. 10.1016/j.pharmthera.2018.03.004.

(22) Curry, A. M.; White, D. S.; Donu, D.; Cen, Y. Human Sirtuin Regulators: The “Success” Stories. Front. Physiol. 2021, 12, 752117. 10.3389/fphys.2021.752117.

(23) Hong, J. Y.; Price, I. R.; Bai, J. J.; Lin, H. A Glycoconjugated SIRT2 Inhibitor with Aqueous Solubility Allows Structure-Based Design of SIRT2 Inhibitors. ACS Chem Biol 2019, 14 (8), 1802– 1810. 10.1021/acschembio.9b00384.

(24) Li, Y.; Liu, T.; Liao, S.; Li, Y.; Lan, Y.; Wang, A.; Wang, Y.; He, B. A Mini-Review on Sirtuin Activity Assays. Biochemical and Biophysical Research Communications 2015, 467 (3), 459–466. 10.1016/j.bbrc.2015.09.172.

(25) Hong, J. Y.; Zhang, X.; Lin, H. HPLC-Based Enzyme Assays for Sirtuins. In ADP-ribosylation and NAD+ Utilizing Enzymes; Chang, P., Ed.; Methods in Molecular Biology; Springer New York: New York, NY, 2018; Vol. 1813, pp 225–234. 10.1007/978-1-4939-8588-3_15.

(26) Chiang, Y.-L.; Lin, H. An Improved Fluorogenic Assay for SIRT1, SIRT2, and SIRT3. Org. Biomol. Chem. 2016, 14 (7), 2186–2190. 10.1039/C5OB02609A.

(27) Wegener, D.; Hildmann, C.; Riester, D.; Schwienhorst, A. Improved Fluorogenic Histone Deacetylase Assay for High-Throughput-Screening Applications. Analytical Biochemistry 2003, 321 (2), 202–208. 10.1016/S0003-2697(03)00426-3.

(28) Wegener, D.; Wirsching, F.; Riester, D.; Schwienhorst, A. A Fluorogenic Histone Deacetylase Assay Well Suited for High-Throughput Activity Screening. Chemistry & Biology 2003, 10 (1), 61– 68. 10.1016/S1074-5521(02)00305-8.

(29) Rossi, A. M.; Taylor, C. W. Analysis of Protein-Ligand Interactions by Fluorescence Polarization. Nat Protoc 2011, 6 (3), 365–387. 10.1038/nprot.2011.305.

(30) Hall, M. D.; Yasgar, A.; Peryea, T.; Braisted, J. C.; Jadhav, A.; Simeonov, A.; Coussens, N. P. Fluorescence Polarization Assays in High-Throughput Screening and Drug Discovery: A Review. Methods Appl. Fluoresc. 2016, 4 (2), 022001. 10.1088/2050-6120/4/2/022001.

(31) Ai, T.; Wilson, D. J.; More, S. S.; Xie, J.; Chen, L. 5-((3-Amidobenzyl)Oxy)Nicotinamides as Sirtuin 2 Inhibitors. J. Med. Chem. 2016, 59 (7), 2928–2941. 10.1021/acs.jmedchem.5b01376.

(32) Borra, M. T.; Langer, M. R.; Slama, J. T.; Denu, J. M. Substrate Specificity and Kinetic Mechanism of the Sir2 Family of NAD^+^ -Dependent Histone/Protein Deacetylases. Biochemistry 2004, 43 (30), 9877–9887. 10.1021/bi049592e.

(33) Avalos, J. L.; Boeke, J. D.; Wolberger, C. Structural Basis for the Mechanism and Regulation of Sir2 Enzymes. Molecular Cell 2004, 13 (5), 639–648. 10.1016/S1097-2765(04)00082-6.

(34) Pan, P. W.; Feldman, J. L.; Devries, M. K.; Dong, A.; Edwards, A. M.; Denu, J. M. Structure and Biochemical Functions of SIRT6. Journal of Biological Chemistry 2011, 286 (16), 14575–14587. 10.1074/jbc.M111.218990.

(35) Feldman, J. L.; Dittenhafer-Reed, K. E.; Kudo, N.; Thelen, J. N.; Ito, A.; Yoshida, M.; Denu, J. M. Kinetic and Structural Basis for Acyl-Group Selectivity and NAD^+^ Dependence in Sirtuin-Catalyzed Deacylation. Biochemistry 2015, 54 (19), 3037–3050. 10.1021/acs.biochem.5b00150.

(36) Hua, L.; Wang, D.; Wang, K.; Wang, Y.; Gu, J.; Zhang, Q.; You, Q.; Wang, L. Design of Tracers in Fluorescence Polarization Assay for Extensive Application in Small Molecule Drug Discovery. J. Med. Chem. 2023, 66 (16), 10934–10958. 10.1021/acs.jmedchem.3c00881.

(37) Zhang, J.-H.; Chung, T. D. Y.; Oldenburg, K. R. A Simple Statistical Parameter for Use in Evaluation and Validation of High Throughput Screening Assays. SLAS Discovery 1999, 4 (2), 67– 73. 10.1177/108705719900400206.

