## Supplementary Figures and synthesis procedure for "A High-throughput Fluorescence Polarization Assay for Screening Sirtuin Inhibitors"

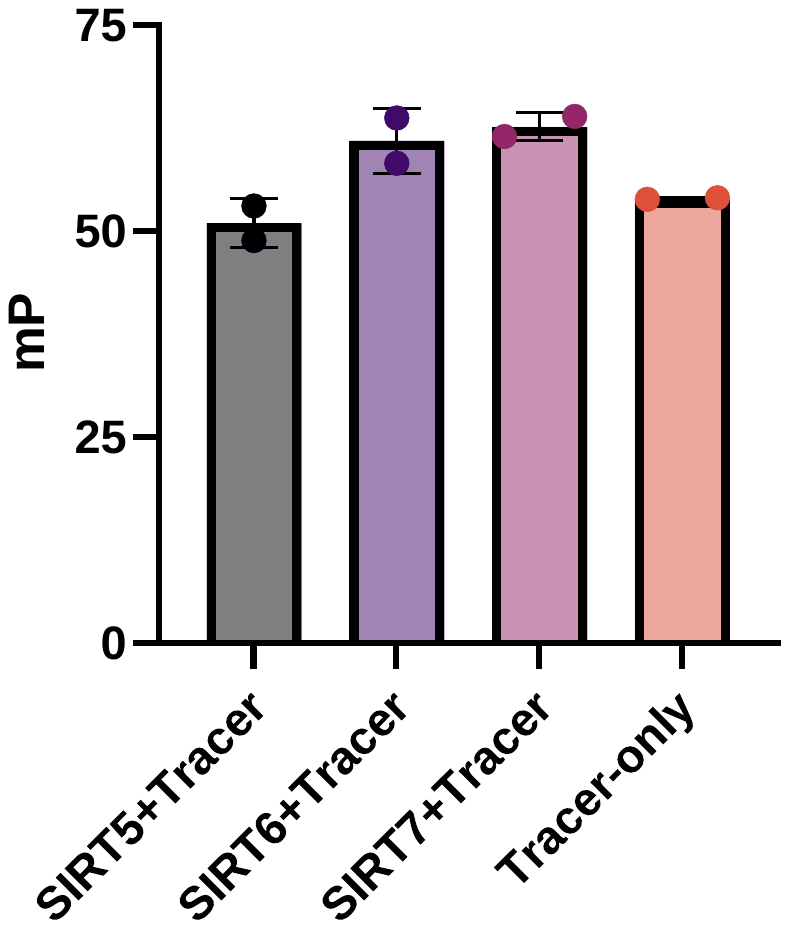

**Figure S1.** Testing the binding of the FP tracer KP-SC-1 for different isoforms of sirtuins – SIRT5, SIRT6, and SIRT7. The protein and tracer concentrations were 500 nM and 2 nM, respectively. Tracer-only control was also included.

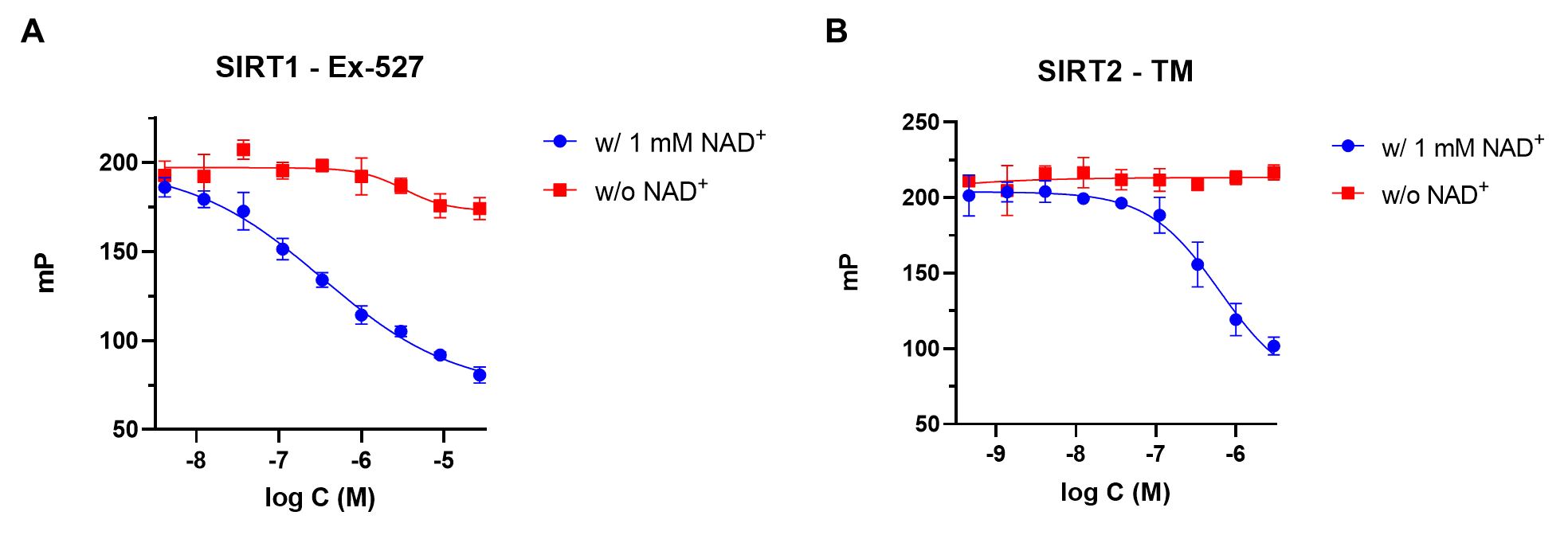

**Figure S2.** In the absence of NAD^+^, Ex-527 (A) and TM (B) do not inhibit SIRT1 and SIRT2, respectively.

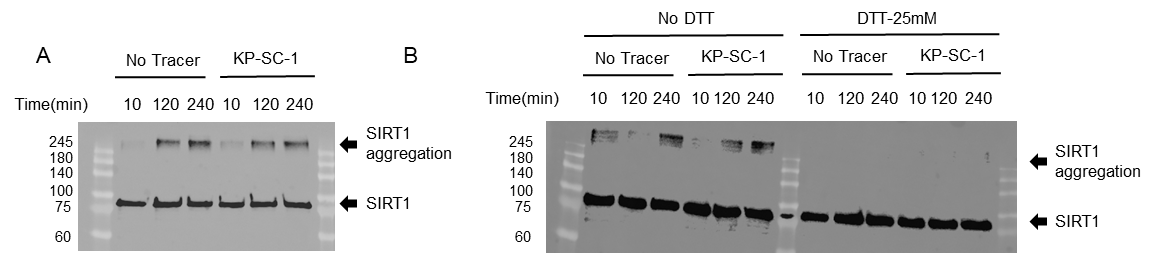

**Figure S3.** Disulfide-bond mediated SIRT1 oligomer formation. (A) Western blot for SIRT1 (60 nM) in the presence or absence of the tracer (5 nM) at indicated incubation times. (B) Western Blot of SIRT1 (60 nM) in the presence or absence of the tracer (5 nM) and DTT at indicated incubation times. The gels were run under reducing conditions.

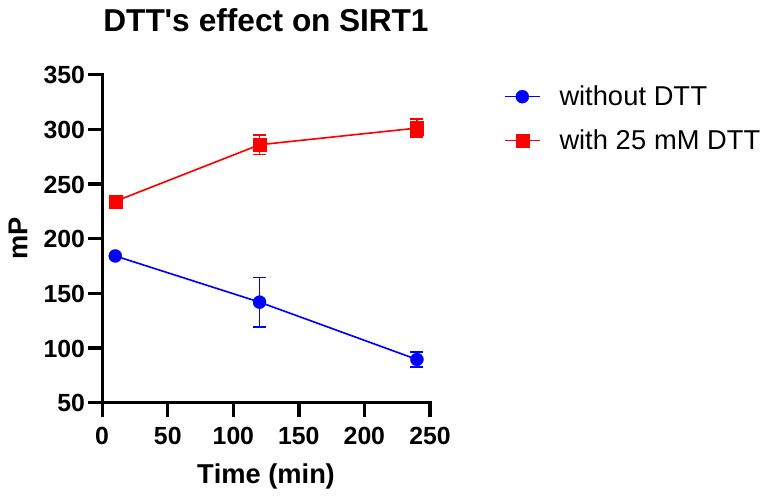

**Figure S4.** The SIRT1 FP assay is stabilized by the addition of DTT.

**Synthesis procedures**

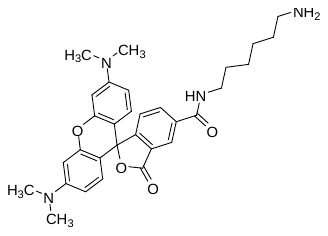

*N-(6-aminohexyl)-3',6'-bis(dimethylamino)-3-oxo-3H-spiro[isobenzofuran-1,9'-xanthene]-5-carboxamide* (**3**)

To a stirring solution of 5-TAMRA (30 mg, 0.070 mmol) and tert-butyl (6-aminohexyl)carbamate (15 mg, 0.070 mmol) in *N,N*-dimethyl formamide (DMF, 1 mL) was sequentially added DIPEA (12.3 μL, 0.070 mmol) and HATU (27 mg, 0.070 mmol). The reaction mixture was allowed to stir for 1 hour before purification on a Combiflash with a C18 column, using water and acetonitrile as solvents. The desired fractions were pooled and lyophilized to afford the pure amide product, which was then dissolved in dichloromethane (DCM, 3 mL) and treated with HCl in dioxane (4 N, 1 mL) overnight. The resulting solution was stirred overnight and directly evaporated under vacuum to afford the hydrochloric acid salt of **3** as a dark purple solid (20 mg, yield: 54%)

**LC-MS (ESI):** m/z calcd for C_31_H_37_N_4_O_4_^+^ [M+H]^+^ 529.3, found 529.2.

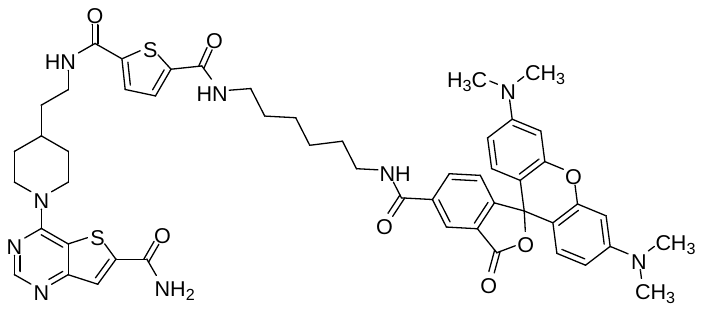

*N^2^-(6-(3',6'-bis(dimethylamino)-3-oxo-3H-spiro[isobenzofuran-1,9'-xanthene]-5-carboxamido)hexyl)-N^5^-(2-(1-(6-carbamoylthieno[3,2-d]pyrimidin-4-yl)piperidin-4-yl)ethyl)thiophene-2,5-dicarboxamide* (KP-SC-1)

Compound **2** (3.4 mg, 7.4 µmol) and **3** (4.7 mg, 8.9 µmol) were dissolved in DMF (500 µL), followed by the addition of DIPEA (3.9 µL, 22.2 µmol). HATU (3.4 mg, 8.9 µmol) was then added, and the reaction was allowed to stand for 1 hour at room temperature. The reaction mixture was directly purified on Combiflash with a C18 column using water and acetonitrile as solvents. Fractions with product were combined and lyophilized to yield pure KP-SC-1 as a dark purple solid (3.1 mg, yield: 36%).

**LC-MS (ESI):** m/z calcd for C_51_H_56_N_9_O_7_S_2_^+^ [M+H]^+^ 970.4, found 970.0.

**^1^H NMR** (400 MHz, DMSO-*d*_6_) δ 8.80 (t, *J* = 5.6 Hz, 1H), 8.61 (d, *J* = 5.7 Hz, 1H), 8.48 (s, 1H), 8.42 (s, 1H), 8.21 (d, *J* = 8.1 Hz, 1H), 8.03 (s, 1H), 7.68 (s, 2H), 7.30 (d, *J* = 8.0 Hz, 1H), 6.50 (s, 5H), 4.69 (d, *J* = 13.2 Hz, 2H), 3.30 (t, *J* = 6.3 Hz, 4H), 3.25 – 3.21 (m, 2H), 3.20 – 3.13 (d, *J* = 8.9 Hz, 5H), 2.94 (s, 12H), 1.89 (d, *J* = 12.9 Hz, 2H), 1.73 (s, 1H), 1.60 – 1.45 (dm, 6H), 1.36 (s, 4H), 1.25 – 1.17 (m, 2H).

**HRMS (ESI+APCI):** m/z calcd for C_51_H_56_N_9_O_7_S_2_^+^ [M+H]^+^ 970.3739, found 970.3758.

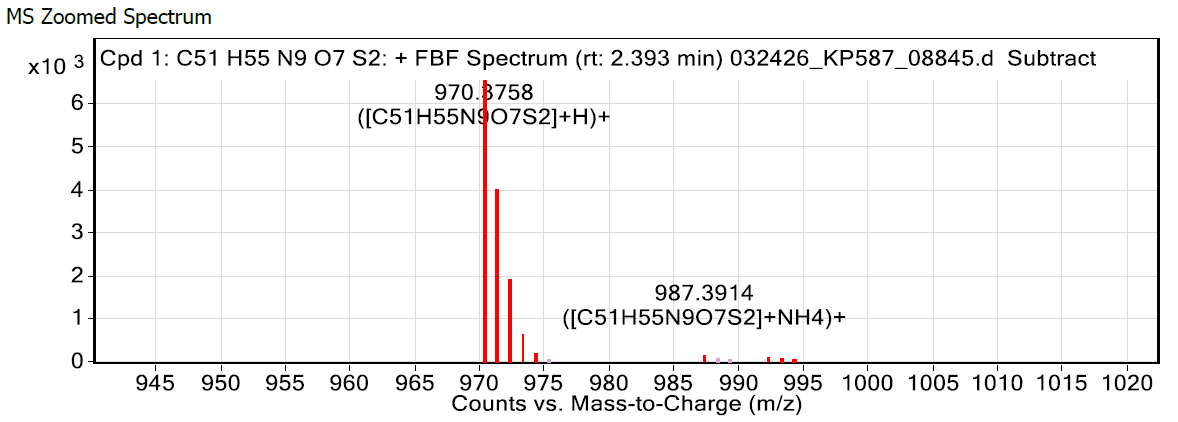

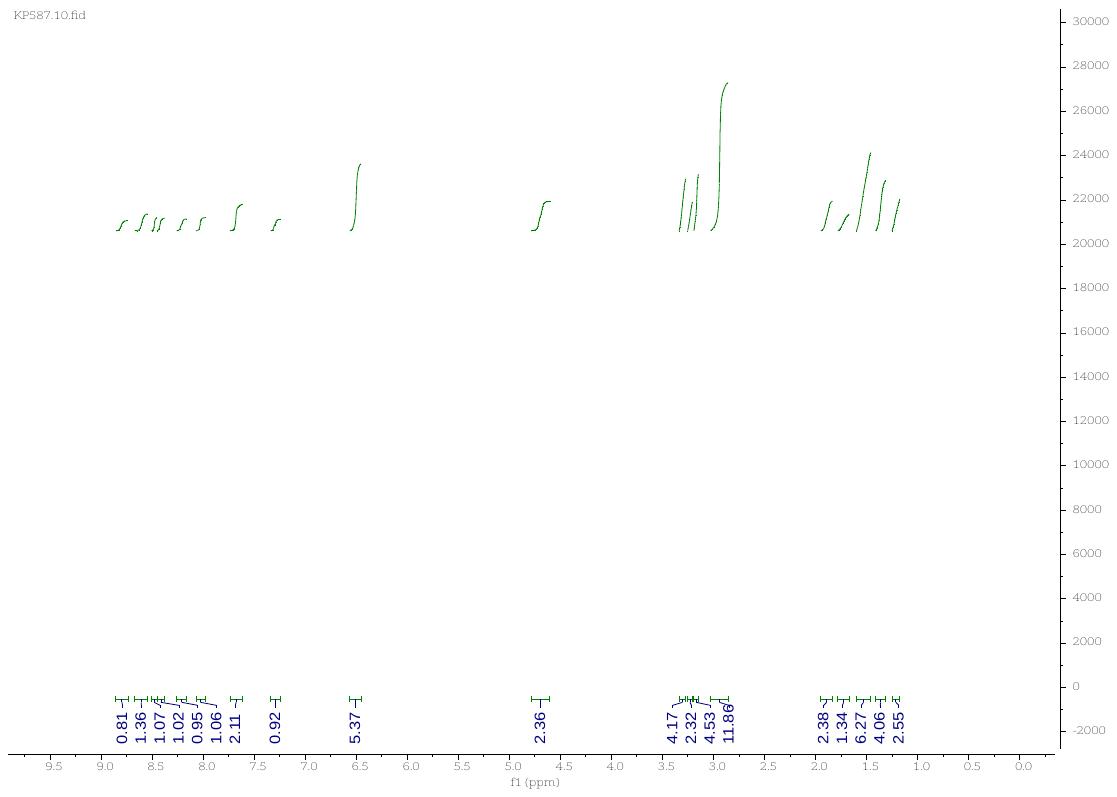
